# Mountain roads across the globe significantly alter local soil microclimates

**DOI:** 10.1101/2024.11.28.625797

**Authors:** Renée Lejeune, Eduardo Fuentes-Lillo, Stef Haesen, Amber Pirée, Dymphna Wiegmans, Lore Hostens, Jonathan Lenoir, Jan Pergl, Michaela Vítková, Tim Seipel, Josef Kutlvašr, Martin A. Nuñez, Romina D. Dimarco, Jake Alexander, Amanda Ratier Backes, Sylvia Haider, Aníbal Pauchard, Ivan Nijs, Jonas J. Lembrechts

## Abstract

Mountain roads have repeatedly shown to host significantly different plant species communities compared to the adjacent natural vegetation. Besides the effect of propagule pressure, altered disturbance regime and soil processes, one of the reasons given for the strong influence of mountain roads on species distributions is a significantly altered microclimate in the roadside compared to the adjacent vegetation, a direct consequence of the altered disturbance regime. However, the microclimatic differences between roadside and natural vegetation have rarely been quantified, particularly lacking global analyses, hampering a better understanding of their importance for mountain biodiversity. In this study, we analysed in-situ measured soil temperatures along mountain roads in seven mountain regions across the globe, in order to assess the impact of mountain roads on a range of bioclimatic variables across the elevational gradient. Our results undeniably show the importance of roadsides as unique microhabitats, even in heterogeneous mountain environments. In most regions, roadside soils had warmer maxima (3.95 ± 2.35°C warmer) and colder minima (0.85 ± 1.11 °C colder) than the soil in the adjacent vegetation, with higher frost risks in winter. Therefore, we recommend future research to incorporate the notion that the local microclimates created by mountain roads could play a critical role in species redistributions in space and time.

## Introduction

Both the effect of mountain roads as well as elevation-dependent temperature changes on plant species communities within high-mountain environments are clear. Plant species communities along mountain roads are known to significantly differ from those in the surrounding natural vegetation (Seipel et al. 2012, Haider et al. 2018). Indeed, both native and non-native species reportedly shift their distributions along mountain roads (Lembrechts et al. 2017, Iseli et al. 2023). These changes in species communities have been linked to a diverse range of factors, including facilitated propagule transportation, disturbance regimes with higher intensity and frequency, altered biotic interactions, and modified soil conditions compared to the adjacent vegetation (Rauschert et al. 2017, Müllerová et al. 2011). Plant species communities along an elevational gradient are expected to experience a higher effect of climate warming with higher latitudes (Mountain Research Initiative EDW Working Group. 2015) and modified microclimatic conditions in mountain roads will enhance this effect even more. However, the nature of the latter, and the magnitude of these microclimatic differences between roadside and natural vegetation, have rarely been quantified (but see e.g. Delgado et al. 2007 for mountain forests) and surely not across multiple sites simultaneously, hampering the importance of their assessment.

Nevertheless, previous research suggests that disturbances similar to those observed in roadsides can significantly alter microclimatic conditions (Delgado et al. 2007). Indeed, soil-level temperatures have been shown to be more extreme (i.e., higher maxima, lower minima) when vegetation cover is reduced due to either natural or human-induced disturbances (Spellerberg 1998, Lembrechts et al. 2018, Haesen et al. 2023). These patterns are well known for lowland forests (De Frenne et al. 2021), yet are likely to be of smaller magnitude in alpine areas, where vegetation cover is often less dense and of shorter stature, which means that disturbance thus causes smaller reductions in said vegetation cover and height (Lembrechts et al. 2014). Additionally, the more complex topography and thus topoclimatic heterogeneity in mountains might obscure these patterns. Finally, microclimate temperatures in mountain soils are, for at least a large part of the year, defined by snow cover (Rixen et al. 2021). Snow cover along roads may be reduced through human practices, for example mechanical snow removal or salt application, as well as through natural processes such as increased solar radiation reaching the ground, or increased wind speeds, which might result in lower winter temperatures and earlier spring onset (Koivusalo and Kokkonen 2002). On the other hand, roadsides might also accumulate snow that is blown off or manually pushed from the road surface towards the road verges (Semádeni-Davies 1999), resulting in more buffered winter temperatures, more humid soil conditions and a later spring onset with altered soil properties (Pomeroy and Brun 2001, Pomeroy et al. 2001). Hence, the expected effect of snow cover on microclimate dynamics at roadsides is not crystal clear due to potential confounding effects.

In this unique multiregional study, we conducted the first-ever analysis of soil temperatures along mountain roads at a large spatial extent, using in-situ measurements from seven mountain regions across the globe. We hypothesize that (1) roadside soil temperatures would be significantly different from those in the adjacent vegetation, with higher maximal and summer soil temperatures. Due to the heterogeneous snow patterns that could be expected, we hypothesize (2) mean and minimum winter soil temperatures in roadsides, dependent on roadside management and regional amount of winter snow, to be higher or lower than in the adjacent vegetation. As a result of the increased maxima and high temperature variability in winter, we anticipate (3) more growing degree days (GDDs) and higher mean annual soil temperatures in roadsides. All effects are expected to be (4) more pronounced at low than at high elevations for maximum temperatures due to the more substantial impact of roadside disturbances on vegetation cover and the negative correlation between elevation and vegetation cover and height (Lembrechts et al. 2014), yet more pronounced at high elevations for minimum temperatures due to a stronger impact of snow at high elevations.

## Methods

### Field monitoring

Temperatures were monitored in the topsoil layer (< 10 cm) of roadsides and adjacent natural vegetation plots, in a systematically paired manner, along mountain roads in seven mountain regions from the Mountain Invasion Research Network (MIREN), across seven countries: Argentina, Chile, Czech Republic, Norway, Spain (Tenerife), Switzerland, and USA (Fig. 1, Haider et al. 2022).

**Figure 1:**
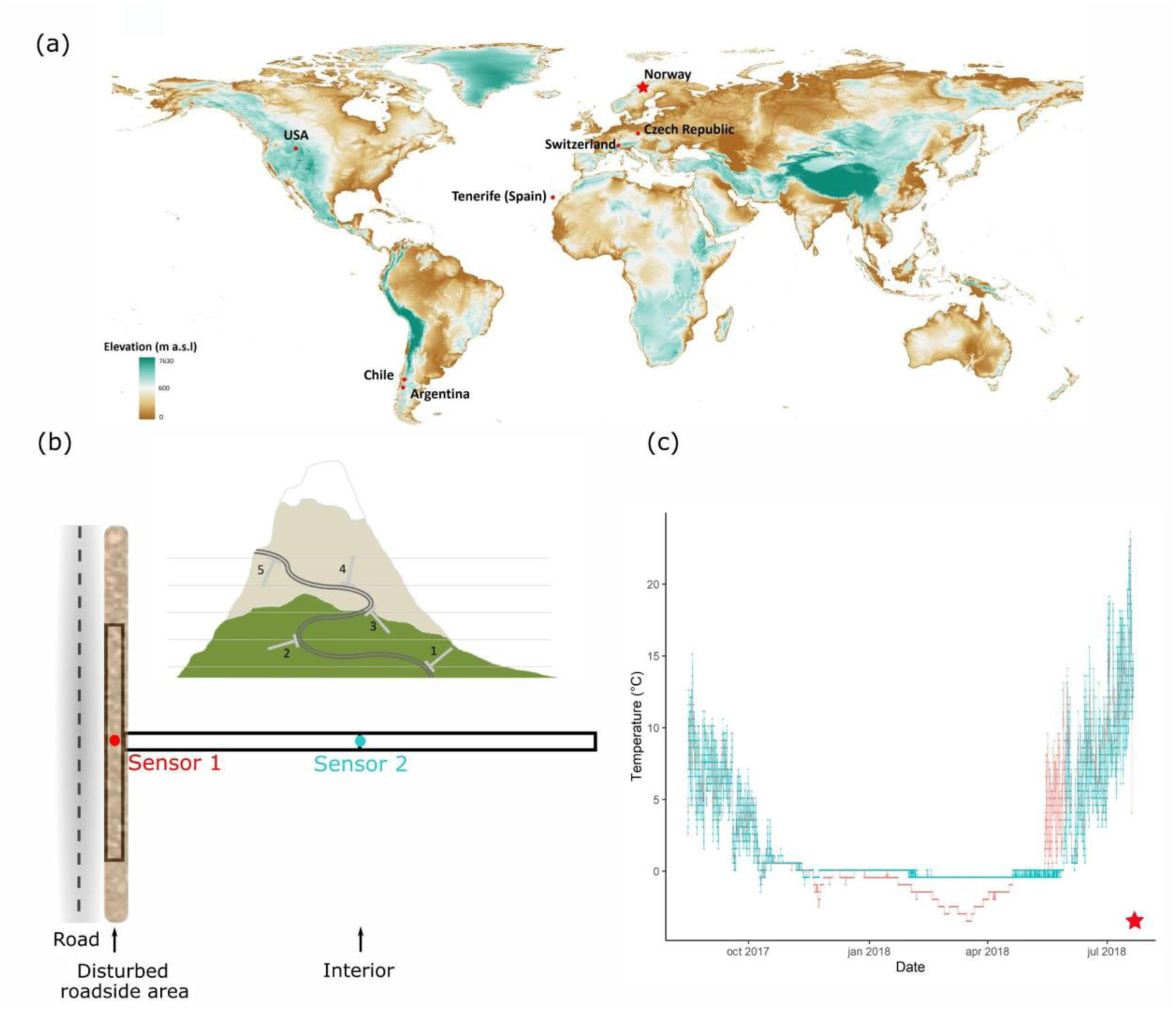
(a) Location of the seven study regions across the globe, on a backdrop of a digital elevation model highlighting high-elevation zones (Tachikawa et al. 2011). Naming convention of regions is following Haider et al. (2022). (b) MIREN Transect design showing the T-shaped transects bordering roadsides along an elevational gradient, as well as the location of the two soil temperature sensors in each transect, in the roadside (sensor 1) and adjacent vegetation (sensor 2). Inset shows the spread of T-transects along a hypothetical mountain road. (c) Example of a one-year temperature time series in a roadside (red) and adjacent vegetation (blue) plot, based on data from a plot at 500 m a.s.l. in Norway (★). Note the quasi-horizontal line during winter in the adjacent vegetation, indicating the insulating effect of snow.

Within the framework of the long-term vegetation monitoring along mountain roads (i.e., paved or gravel roads open to motorized vehicles) directed by the MIREN network (www.mountaininvasions.org, Haider et al. 2022), each region selected up to three sample roads that extend over a broad elevational gradient, from the bottom of the mountain region, in a valley, at sea level, or where no further elevation change occurred, up till the highest elevation reached by roads in the region. The elevation gradient of each road was then divided into 19 equally wide elevational bands from the lowest to the highest possible sampling location, resulting in a total of 20 sample sites per road. At each of these sample sites, three 2 m x 50 m vegetation survey plots were laid out in the form of a “T”-shaped transect, one plot parallel to the road and the other two extending end-to-end and perpendicular to the road, starting from the center of the first plot, with midpoints at 25 m and 75 m from the roadside plot (Fig. 1b, Haider et al. 2022).

In the seven regions of interest, these T-shaped vegetation plots have been installed at various moments since 2006 and are aimed for repeated monitoring every five years. In all or a selection of these vegetation plots, small, rugged temperature sensors (iButtons DS1921G or DS1922L (Maxim Integrated), HOBO Pendant UA-001-08 (Onset) or TOMST TMS4s (shortened; 23 cm instead of the standard 29 cm; Wild et al. 2019, Table 1)) were installed, each time pairwise: one sensor in the middle of the roadside plot vs. one at the start of the second perpendicular plot (i.e., at 50 m from the roadside plot). iButtons were wrapped in parafilm to prevent moisture damage and put into a small Ziplock bag; iButtons and HOBOs were buried just below the soil surface (0-5 cm). The shortened TOMST TMS4s sensors were installed vertically, with their top at the soil surface, and the data from the second sensor (6 cm below the soil surface) was used. Note that when measuring the temperature of soil below the surface, no radiative heat is supplied to a thermometer and the errors in measurement resulting from e.g., sensor brand or wrapping are likely to be negligible (Maclean et al. 2021). Sensors measured temperatures every 15 to 240 minutes over a period of at least 12 months between 2014 and 2022 (Table 1, Fig. 1). In all regions except Tenerife (Spain), Czech Republic and Argentina, plot-level forest cover was documented over the elevational gradient as percentage covered by trees per T-transect.

**Table 1:**
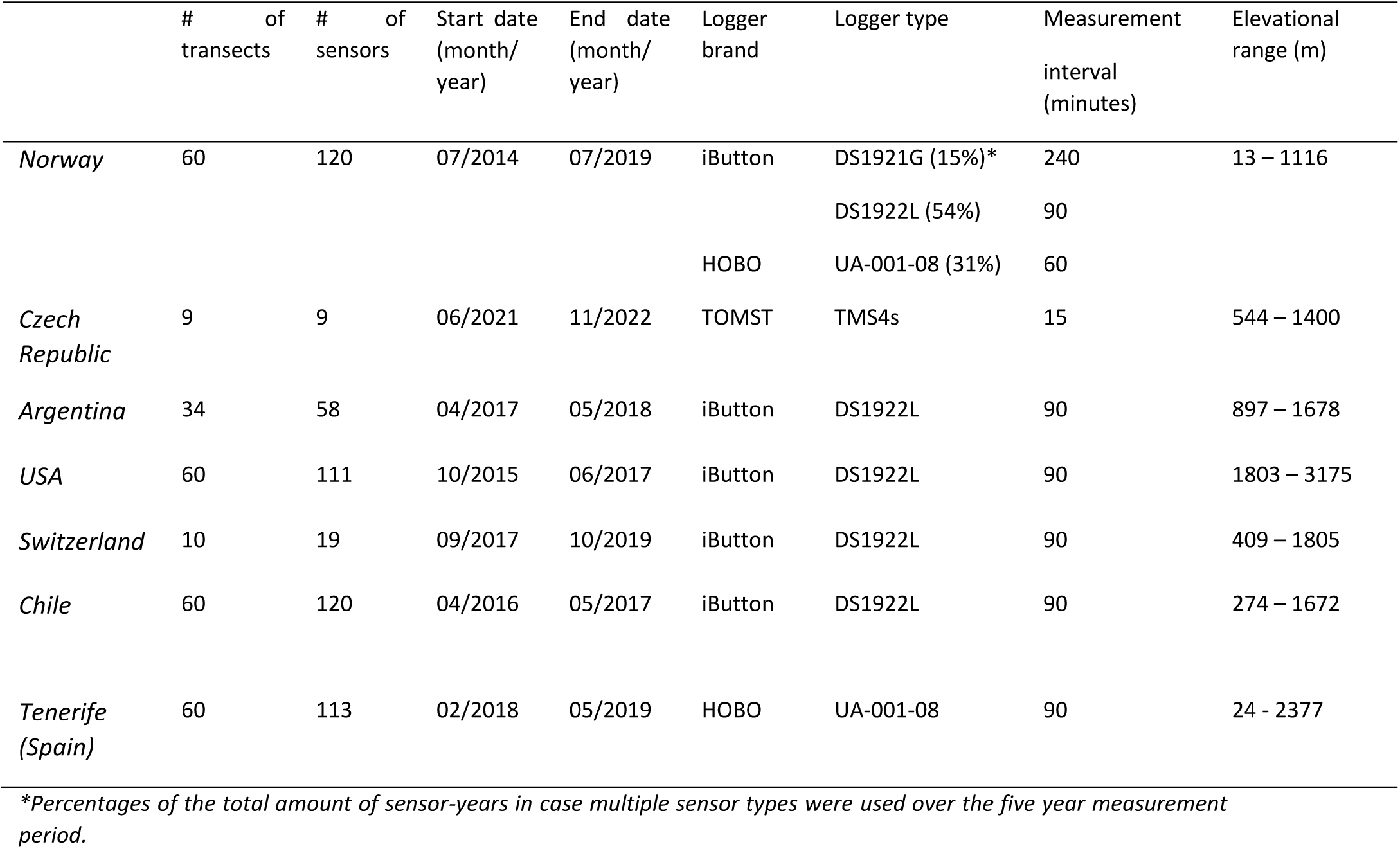
Number of transects, number of sensors, start date and end date of soil temperature measurements, logger brand and type, measurement interval, and elevational range for all seven mountain regions. Regions here and throughout are ordered from low to high mean annual soil temperature in their lowest elevation plot.

### Bioclimatic variables

The abovementioned sensor data was used to calculate a series of bioclimatic variables per sensor: mean annual temperature (averages over a 12-month period; BIO1 following the ANUCLIM framework, Xu & Hutchinson 2011), mean summer (BIO10) and winter (BIO11) temperature (average over respectively July till September for summer and December till February for winter on the northern hemisphere, and vice versa on the southern hemisphere, as these were in each region identified as the warmest and coldest quarter based on the available data), growing degree days (GDD, sum of all daily mean temperatures in a year over a certain threshold, with the threshold set here at 0°C; the only variable not following the ANUCLIM framework), and the maximum (BIO5) and minimum (BIO6) temperature of respectively the warmest and coldest month of the year. BIO5 was calculated as the 5% highest daily maximum temperature in a month, to remove extreme outliers. Similarly, BIO6 was defined as the 5% lowest daily minimum in a month.

### Statistical analysis

All analyses were performed in R version 4.2.1. Each bioclimatic variable was modeled as a function of plot-level elevation (in m a.s.l.), distance to the road (binary: roadside vs. 50 m distant from the roadside plot in adjacent vegetation) and their interactions. Additionally, each bioclimatic variable was modeled as a function of forest cover of the adjacent plot (in percentage), distance to the road (see earlier) and their interactions in separate models due to the correlation between elevation and forest cover (see Appendix A). Since both roadside and adjacent plots within one transect received the same forest cover value, this model only accounts for the change in forest cover over elevation. Models were made using linear mixed models (package *lme4,* Bates et al. 2015), with region as a random intercept and elevation times distance to the road as a random slope. This way, we could obtain region-specific coefficients for both the elevation and road parameter and their interaction using partial pooling (Harrisson et al. 2018). As temperature differences tend to be smaller when approaching 0°C, bioclimatic variables were log-transformed. To avoid log-transforming the few negative temperature values, we rescaled temperature values by subtracting the lowest negative temperature value from all of them, rendering them all positive. Using the same data, an assessment of global emerging patterns was made for each bioclimatic variable to visualize the effect of both elevation and forest cover independent of region. For this a reduced version of the above described model was used and elevation was scaled as the different regions had various elevational ranges. Model predictions were visualized using the *ggplot2* package (Wickham 2011).

## Results

### Regional trends

At the regional scale, soil temperature patterns in roadsides broadly reflected those observed in the global model, with notable regional variations (Fig. 2-3; Tables 2 & 3). Mean annual soil temperatures were consistently higher in roadsides than adjacent vegetation in six out of seven regions or became higher at higher elevations in one region. Roadside temperatures were, on average, 0.58 ± 0.66°C higher in lowlands and 0.51 ± 0.33°C higher in highlands. Mean summer temperatures followed a similar trend, being consistently higher in roadsides across all regions (1.30 ± 1.41°C). Mean winter soil temperatures, however, were lower in roadsides in three regions and became lower with elevation in four regions, averaging 0.33 ± 0.73°C lower than in adjacent vegetation (Fig. 2; Table 2).

**Figure 2:**
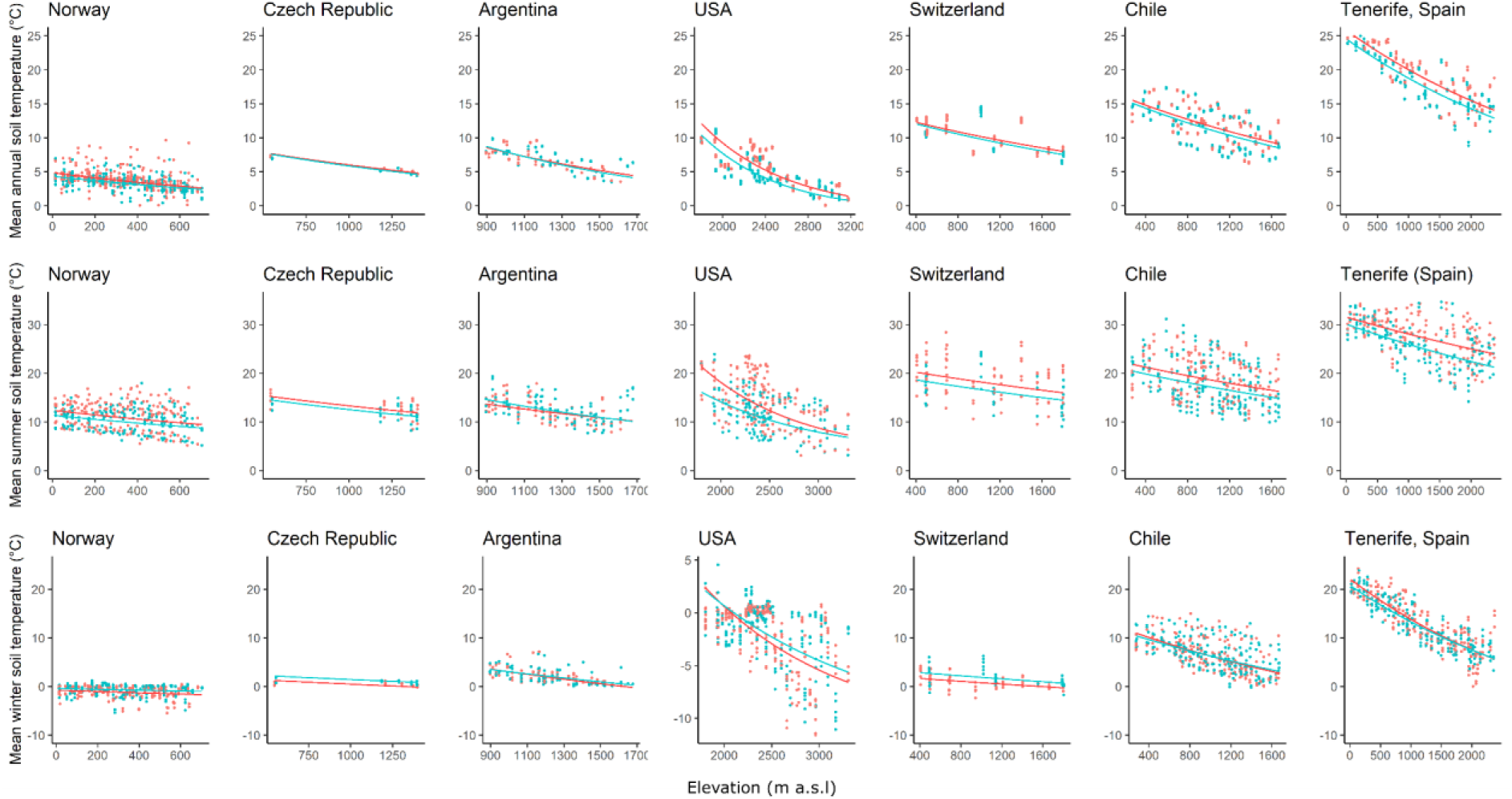
Key soil bioclimatic variables as a function of elevation (x-axis) and distance to the road (roadside = red, natural vegetation = blue) for seven mountain regions. Top row: mean annual soil temperature. Middle row: monthly mean soil temperature during summer (July-September on the northern hemisphere, December-February on the southern hemisphere). Bottom row: monthly mean soil temperature during winter (December-February on the northern hemisphere, July-September on the southern hemisphere). Points are raw measurements, fitted curves are from a partial pooling procedure on linear mixed models using the logarithm of the dependent variable (see Table 1). Regions ordered from lowest to highest mean annual temperature in their lowest elevation plot.

**Figure 3:**
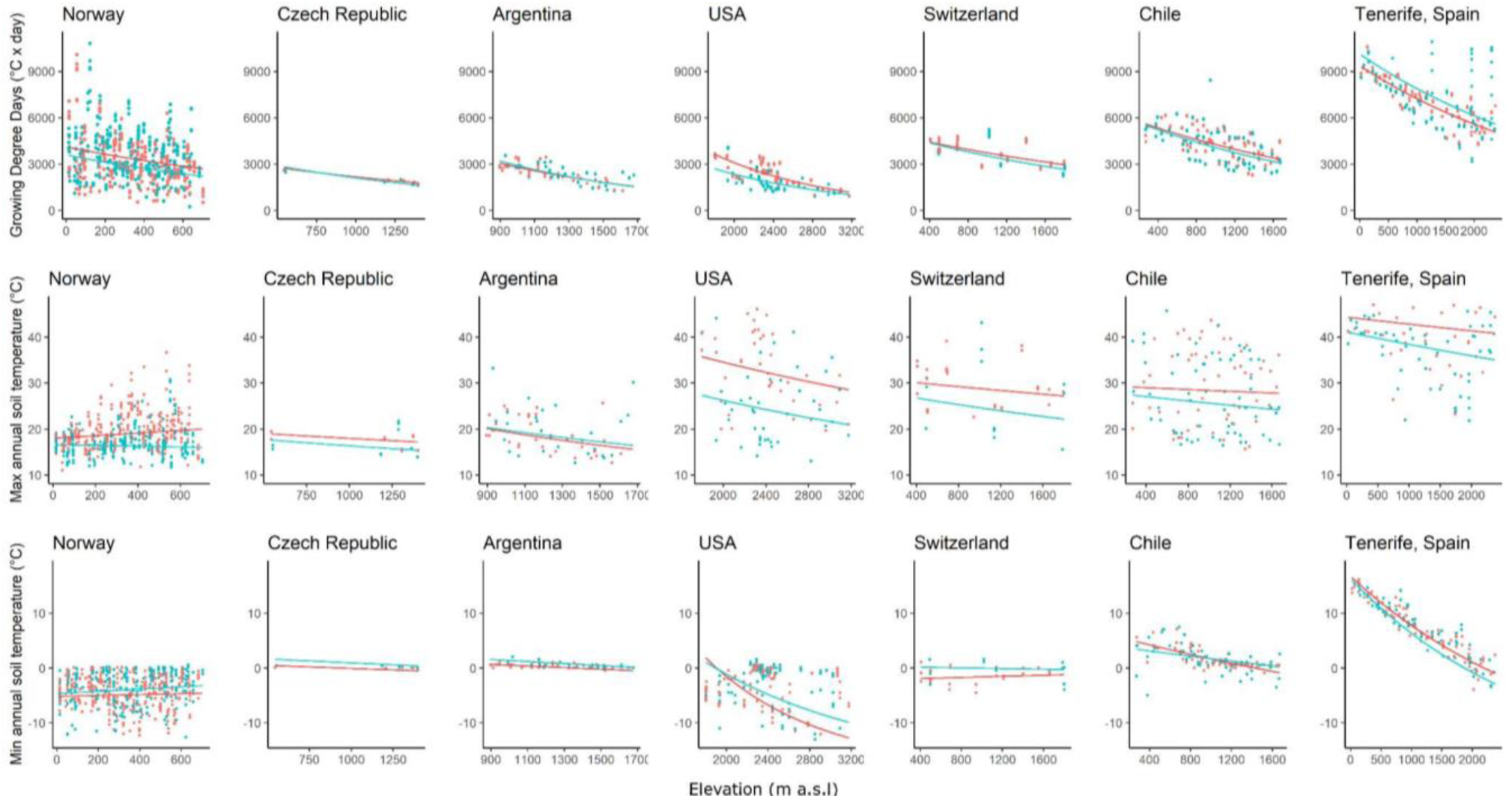
Key soil bioclimatic variables as a function of elevation (x-axis) and distance to the road (roadside = red, natural vegetation = blue) for seven mountain regions. Top row: Growing Degree Days (GDDs). Middle row: annual maximum soil temperature. Bottom row: annual minimum soil temperature. Points are raw measurements, fitted curves are from a partial pooling exercise on linear mixed models using the logarithm of the dependent variable (see Table 2). Regions are ordered from lowest to highest mean annual temperature in their lowest elevation plot.

**Table 2:** (a) Global coefficients (including p-values) and region-specific coefficients for key soil bioclimatic variables as a function of elevation and distance to the road for seven mountain regions, as in Fig. 2. Top row: mean annual soil temperature. Middle row: monthly mean soil temperature during summer (July-September on the northern hemisphere, December-February on the southern hemisphere). Bottom row: monthly mean soil temperature during winter (December-February on the northern hemisphere, July-September on the southern hemisphere). Coefficients are derived from a partial pooling procedure on linear mixed models using the logarithm of the dependent variable. Positive coefficients in black, negative coefficients in red. (b) Modeled temperature differences between the roadside and the adjacent at the lowest (lowland) and highest (highland) elevation of each region’s elevational gradient. Negative differences (roadsides cooler than adjacent) in red. Regions are ordered from lowest to highest mean annual temperature in their lowest elevation plot.

| (a) |  |  |  |  | (b) |  |
| --- | --- | --- | --- | --- | --- | --- |
|  | Intercept | Elevation | Road | Elevation * Road | Roadside T relative to adjacent |  |
| Mean annual soil temperature (°C) |  |  |  |  | Lowland | Highland |
| Global | 2.81<br>(**<0.001**) | **-0.00062**<br>(**<0.001**) | 0.0028<br>(0.92) | 0.000089<br>(**<0.001**) | +0.92 | +0.79 |
| Norway | 1.66 | **-0.00065** | 0.097 | **-0.000046** | +0.53 | +0.22 |
| Czech Republic | 2.41 | **-0.00050** | **-0.0052** | 0.000029 | +0.089 | +0.20 |
| Argentina | 3.01 | **-0.00083** | **-0.11** | 0.000099 | **-0.16** | +0.32 |
| USA | 4.88 | **-0.0014** | **-0.079** | 0.00012 | +1.69 | +0.62 |
| Switzerland | 2.69 | **-0.00031** | 0.0062 | 0.000027 | +0.23 | +0.48 |
| Chile | 2.88 | **-0.00038** | 0.020 | 0.000023 | +0.43 | +0.57 |
| Tenerife (Spain) | 3.24 | **-0.00026** | 0.048 | 0.000014 | +1.26 | +1.15 |
| Mean summer soil temperature (°C) |  |  |  |  |  |  |
| Global | 2.78<br>(**<0.001**) | **-0.0003**<br>(**<0.001**) | 0.057<br>(0.096) | 0.000052 (0.030) | +1.71 | +1.02 |
| Norway | 2.12 | **-0.00054** | 0.11 | 0.0000077 | +0.99 | +0.72 |
| Czech Republic | 2.66 | **-0.00041** | 0.041 | 0.000037 | +0.72 | +0.78 |
| Argentina | 3.01 | **-0.00063** | **-0.20** | 0.00013 | **-0.94** | +0.10 |
| USA | 4.06 | **-0.00083** | 0.60 | **-0.00014** | +5.24 | +0.48 |
| Switzerland | 2.84 | **-0.00022** | 0.088 | 0.000021 | +1.57 | +1.52 |
| Chile | 2.94 | **-0.00028** | 0.067 | 0.000029 | +1.37 | +1.45 |
| Tenerife (Spain) | 3.30 | **-0.00017** | 0.051 | 0.000037 | +1.45 | +2.74 |
| Mean winter soil temperature (°C) |  |  |  |  |  |  |
| **Global** | **3.00**<br>**(<0.001)** | **-0.0003**<br>**(<0.001)** | **-0.0055**<br>**(0.82)** | **-0.000017**<br>**(0.25)** | **+0.18** | **-0.79** |
| Norway | 2.42 | -0.000077 | -0.034 | -0.000047 | -0.38 | -0.69 |
| Czech Republic | 2.68 | -0.00012 | -0.061 | -0.000014 | -0.92 | -0.97 |
| Argentina | 2.99 | -0.00031 | 0.063 | -0.000064 | +0.08 | -0.51 |
| USA | 3.61 | -0.00055 | 0.26 | -0.00013 | +0.28 | -0.97 |
| Switzerland | 2.72 | -0.00012 | -0.088 | 0.0000032 | -1.21 | -0.98 |
| Chile | 3.17 | -0.00029 | 0.036 | -0.000037 | +0.58 | -0.37 |
| Tenerife (Spain) | 3.48 | -0.00026 | 0.041 | -0.000019 | +1.33 | -0.088 |

Similarly, growing degree days (GDD) were higher in roadsides than adjacent vegetation in three regions and increased with elevation in another three, averaging 187.92 ± 320.30°C·days higher across these regions (Fig. 3; Table 3). Annual temperature maxima were consistently higher in roadsides in six regions with on average 3.95 ± 2.35°C, while annual minima were consistently lower in three regions or became lower at higher elevations in three others (0.85 ± 1.11 °C lower on average). Importantly, while adjacent vegetation temperatures declined with elevation, roadside temperatures sometimes increased, such as for annual maxima in Norway and minima in Switzerland. The USA showed the largest roadside-to-vegetation difference for annual maximum soil temperatures (Table 2).

**Table 3:**
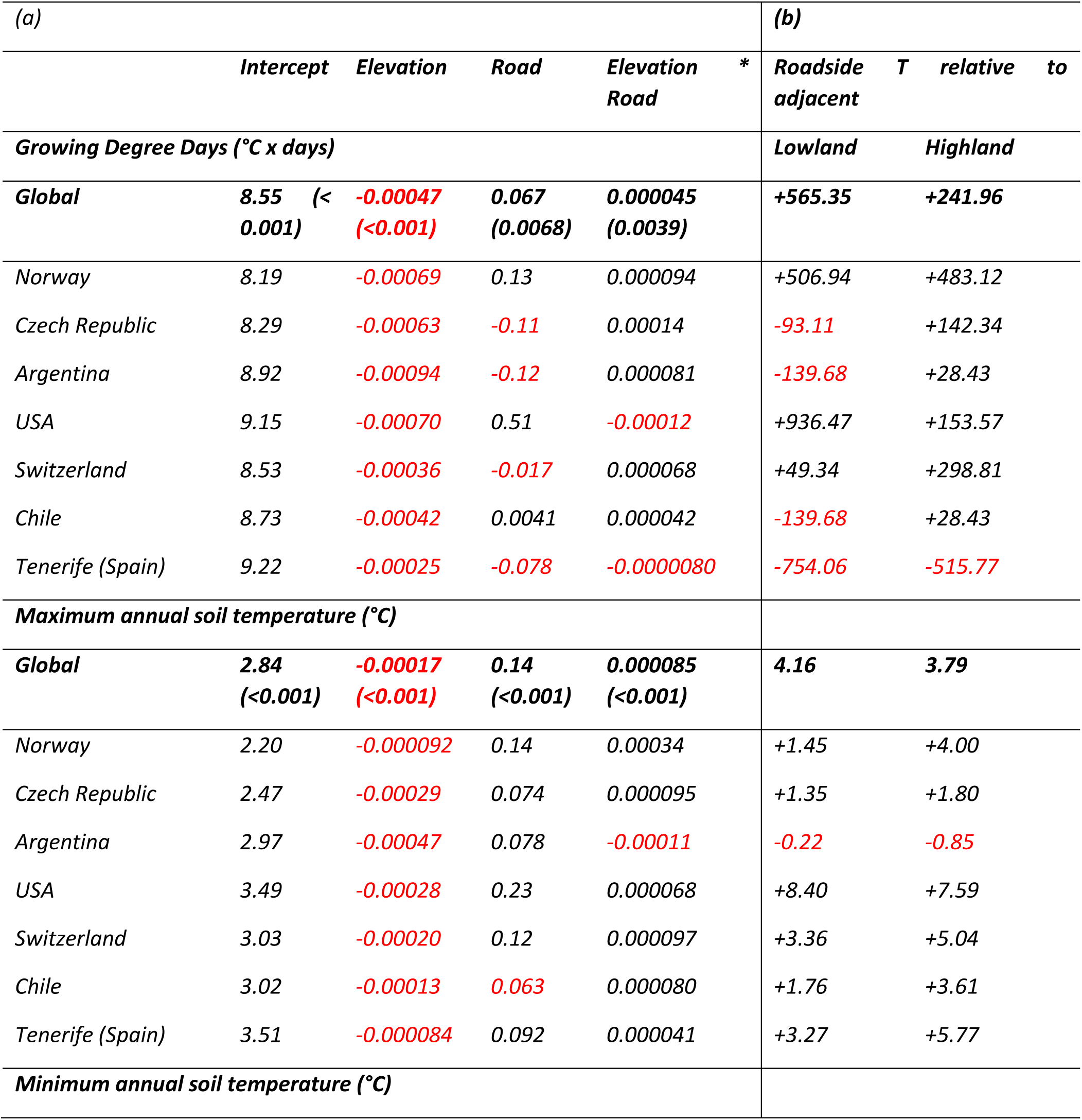
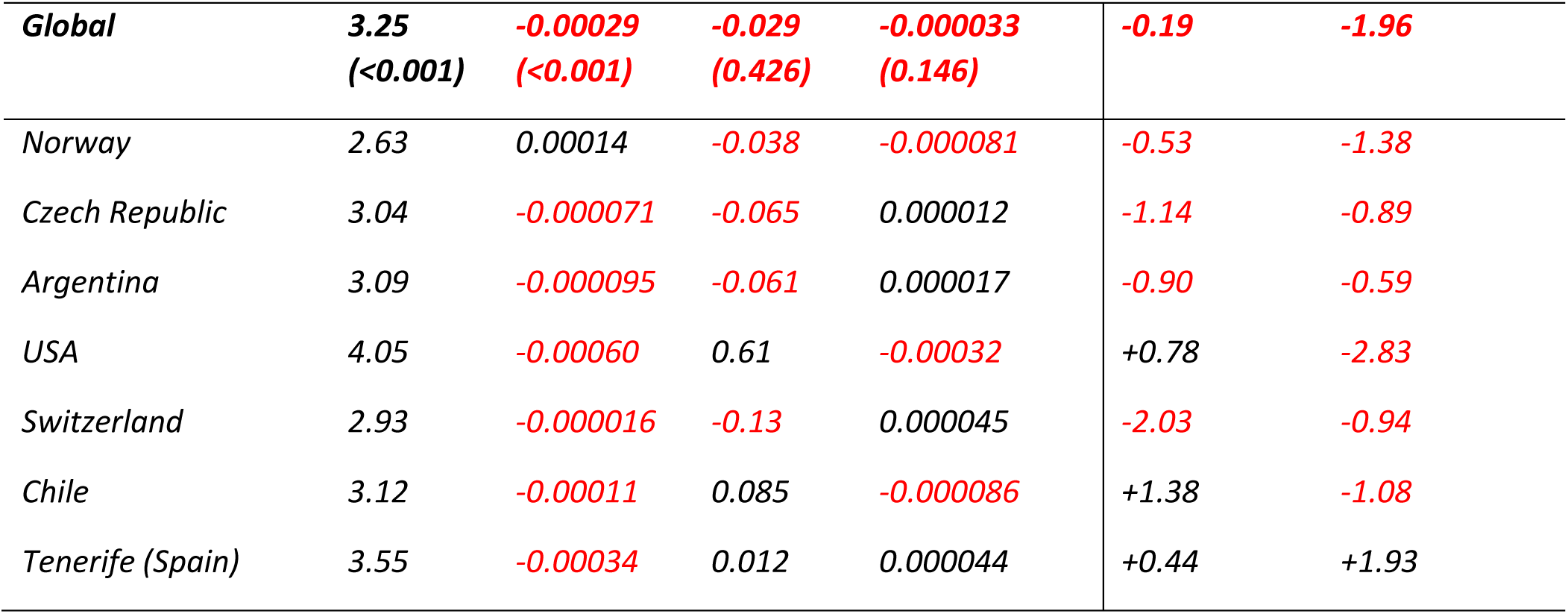
(a) Global coefficients (including p-values) and region-specific coefficients for key soil bioclimatic variables as a function of elevation (m a.s.l.) and distance to the road (road versus adjacent) for seven mountain regions, as in Fig. 3. Top row: Growing Degree Days. Middle row: annual maximum soil temperature (95% highest). Bottom row: annual minimum soil temperature (5% lowest). Coefficients are derived from a partial pooling exercise on linear mixed models using the logarithm of the dependent variable. Positive coefficients in black, negative coefficients in red. (b) Modeled temperature differences between the roadside and the adjacent at the lowest (lowland) and highest (highland) elevation of each region’s elevational gradient. Negative differences (roadsides cooler than adjacent) in red. Regions are ordered from lowest to highest mean annual temperature in their lowest elevation plot.

The effect of forest cover, though not significant in all regions, revealed distinct trends where present. Mean annual and summer soil temperatures increased with forest cover in half of the regions (warming effect) but decreased in the other half (cooling effect, Appendix fig. A3; Appendix table A1). Roadside temperatures consistently exceeded those in adjacent vegetation across all regions. GDD and annual maximum soil temperatures typically decreased with forest cover but were consistently higher in roadsides. Conversely, annual minimum soil temperatures were lower in roadsides or declined further with forest cover in all regions, though they increased with forest cover in most cases, except in Chile (Appendix table A2; Appendix fig. A1). As with elevation, the USA showed the largest roadside-to-vegetation difference for annual maximum soil temperatures (Appendix table A2).

### Global model

Across all regions, mean annual and mean summer temperatures, growing degree days, and maximum annual soil temperatures were higher in roadsides compared to adjacent vegetation (Fig. 4; Tables 2 & 3). Conversely, annual minimum and mean winter soil temperatures were lower in roadsides. Elevation significantly influenced all temperature variables, often interacting with road disturbance. For instance, mean winter temperatures in roadsides were 1°C cooler than in adjacent vegetation at high elevations but only 0.18°C cooler at low elevations (Table 2).

**Figure 4:**
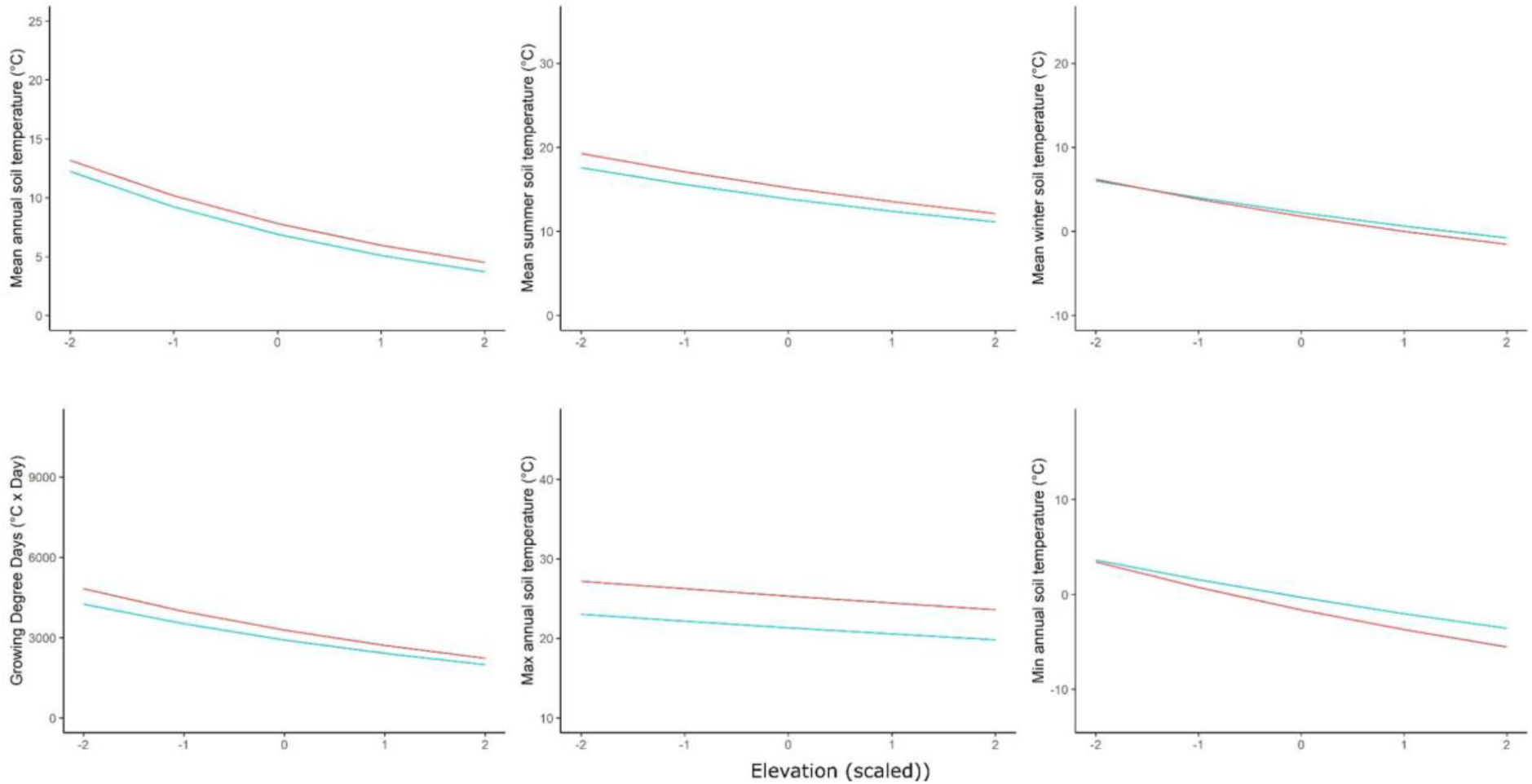
Key soil bioclimatic variables as a function of elevation (x-axis, elevational values scaled within each region with a mean of zero and a standard deviation of one) and distance to the road (roadside = red, natural vegetation = blue) across all seven mountain regions. Top row: mean annual soil temperature, monthly mean soil temperature during summer (July-September on the northern hemisphere, December-February on the southern hemisphere) and monthly mean soil temperature during winter (December-February on the northern hemisphere, July-September on the southern hemisphere). Bottom row: Growing Degree Days (GDDs), annual maximum soil temperature and annual minimum soil temperature. Fitted curves are drawn using the coefficients of the fixed effects of linear mixed models using the logarithm of the dependent variable (see Table 1).

The effect of forest cover on climatic variables was significant for mean summer, minimum annual, and maximum annual soil temperatures. Most notably, for all variables, there was a positive interaction between distance to the road and forest cover: temperatures in roadsides were substantially higher relative to adjacent vegetation at sites with high forest cover compared to those with low forest cover (Appendix tables A1 & A2, Appendix Fig. A3).

## Discussion

Microclimates in mountain roadsides were highly different – often surpassing a temperature difference of 1°C – in comparison to those found in the adjacent natural vegetation. In fact, the soil temperatures observed in roadside locations were frequently more similar to those in adjacent vegetation plots hundreds of meters up or down the elevational gradient, rather than temperatures only 50 m further in the adjacent vegetation. Indeed, our models showed an average elevational lapse rate of 0.45°C/100 m in the natural vegetation, smaller than this 1°C difference (note that the average global macroclimatic lapse rate is estimated at 0.65°C/100 m; ICAO 1993). While we observed significant variation in these trends between regions, we especially found strong evidence for warmer annual maxima and summer mean soil temperatures along roadsides compared to the adjacent vegetation, and lower annual minima and winter temperatures in roadsides at high elevations. Looking at the forest cover gradient the same patterns for annual maxima and summer soil temperatures, warmer along roadsides compared to the adjacent vegetation, were found . These findings suggest a greater coupling between macro- and microclimatic fluctuations along roadsides than in the adjacent vegetation, implying that the natural vegetation acts as a buffer against changes in macroclimatic conditions, though a potential amplifying effect of roadsides in comparison to temperature fluctuations as measured by regular weather stations cannot be ruled out.

As a result of the above, mean annual temperatures and GDDs were often higher in roadsides than in the adjacent vegetation, the increase in GDD being even more pronounced when modeled against forest cover. This implies that there is a greater buffering effect with higher forest cover, and thus a larger warming in the roadside. Roadside microclimatic conditions could therefore be beneficial to species coming from warmer, lower elevation areas, in line with the observed significant upward shift in elevational distributions for both native and non-native lowland species along the roads in several of the studied mountain regions (Lembrechts et al. 2017, Iseli et al. 2023). While it is unlikely that these altered microhabitat conditions are the only determinant of these elevational shifts, it has been shown experimentally that locally increased GDDs can significantly improve establishment success of non-native plant species above their current elevational distributions (Lembrechts et al. 2016, Lembrechts et al. 2017).

As expected, roadside effects were more consistent for mean, summer, and maximum temperatures than for winter temperatures and minima. The latter temperature differences indeed result at least partially from differences in snow cover among regions and between roadside and adjacent vegetation, with roadside snow cover likely to be highly locally heterogeneous within mountain regions where part of the elevation gradient is covered by snow during a significant period of time. Indeed, the main roads are often cleared of snow, yet roadsides might accumulate additional snow due to wind patterns and mechanical accumulation (Semádeni-Davies 1999). Additionally, snow clearing might be limited to lower elevation sites. Despite this complexity, winter temperatures and minima were shown to be more often lower in roadsides than in the adjacent vegetation, suggesting that snow cover loss is a more important process than snow accumulation, at least along the studied roads. This would be in line with the often observed higher GDDs along roadsides, which could at least partially result from earlier snowmelt in spring (Wipf et al. 2006). This important role of snow is supported by the fact that in regions with the coldest climates, such as Norway and the USA, winter temperatures were consistently lower in the roadside than in the adjacent vegetation. That pattern was clearly reversed in the warmer climate of Tenerife, especially at low elevations, where snow did not occur.

The observed lower minima and winter temperatures in roadsides compared to the adjacent vegetation could at first glance be considered the drivers of the observed downward shifts in high elevation species along mountain roads (Lembrechts et al. 2017). However, most high elevation species are not limited by climate at the trailing edge of their distribution, yet by biotic interactions with more competitive lowland species (Boulangeat et al. 2012). It is thus more likely that the reduced vegetation cover in mountain roads, and the increased propagule transport along the roadside corridor are driving these downward shifts (Lembrechts et al. 2017). Additionally, the lower minima and thus increased frost risks in roadside soils did not stop the observed upward expansion of lowland species, suggesting that the average 1.30°C increase in summer temperatures and 187.92 °C x days increase in GDD outweighed the 0.33°C drop in mean winter temperatures.

Interestingly, local adiabatic lapse rates were often less steep for mean annual, mean summer, and annual maximum temperatures in the roadside than in the natural vegetation, resulting in higher positive offsets in the roadsides at high than at low elevations. This was contrary to our expectations, yet might result from a strong shading at both low and high elevation in the adjacent vegetation while in the roadside there is strong shading only at low elevation. At high elevations, however, the adjacent vegetation is often too small to shade disturbed roadsides, and its shading effect is thus limited to the soil growing right underneath it, so the intrinsically colder (i.e. high-elevation) sites would warm more (in summer). This would reduce the temperature gradient, hence the lapse rate. For mean winter temperatures, differences between roadside and adjacent vegetation were as expected larger at high elevations, potentially due to larger differences in snow cover.

Patterns were largely but not entirely consistent between regions, showing that the impact of roads on microclimate cannot simply be generalized across the globe. Part of these differences could be attributed to macroclimatic conditions in the region, as already shown above for mean winter temperatures, but can also be caused by differences in roadside management, such as the mowing regime, between regions. Similarly, the exceptionally low mean summer temperatures in roadsides in Argentina, especially at low elevations, could have resulted from the planting of trees in the roadsides, in a landscape otherwise dominated by short-statured grasses. However, the lack of forest cover data in Argentia creates uncertainty in this conclusion.

## Conclusions and implications

Our results undeniably show the importance of roadsides creating different microclimate conditions in heterogeneous mountain environments. In most regions, roadside soils had warmer and colder extremes than the adjacent vegetation, with lower cold extremes in winter and higher risks of extreme heat during summer, but also longer growing seasons in summer. These temperature trends could potentially explain some of the observed substantial upward shifts in lowland native and non-native species distributions along mountain roads (Lembrechts et al. 2016).

We therefore recommend future research to incorporate the notion that local microclimates created by disturbance could play a critical role in species redistributions in space and time and further enhance effects of climate change. Our results show that these disturbance effects can act on a scale of 50 meters (and likely less). Incorporating these disturbance effects on the microclimate in large-scale assessments of species distributions will require high-resolution gridded microclimate products, created by models that include land use and land use change in their assessments (Lembrechts and Nijs, 2020, De Frenne et al. 2021).

## Acknowledgements

This research has been funded by the Research Foundation Flanders (12P1819N, G018919N and 1512720N) and BiodivERsA+-projects ASICS (G0H6720N, BiodivERsA+, BiodivClim call 2019–2020) and Forest-Web-3.0 (G0GDZ23N, BiodivERsA+, BiodivClim call 2023). AP and EFL acknowledge funding by Fondecyt 1180205, Fondecyt 1231616, ART 210038 and ANID/BASAL FB210006. SH was supported by a FLOF fellowship (project nr. 3E190655) of the KU Leuven. MV, JP and JK gratefully acknowledge the support of BiodivClim Call 2019 (TACR SS70010001) and long-term research development project RVO 67985939 (Czech Academy of Sciences).

## Appendix A

**Figure A1:**
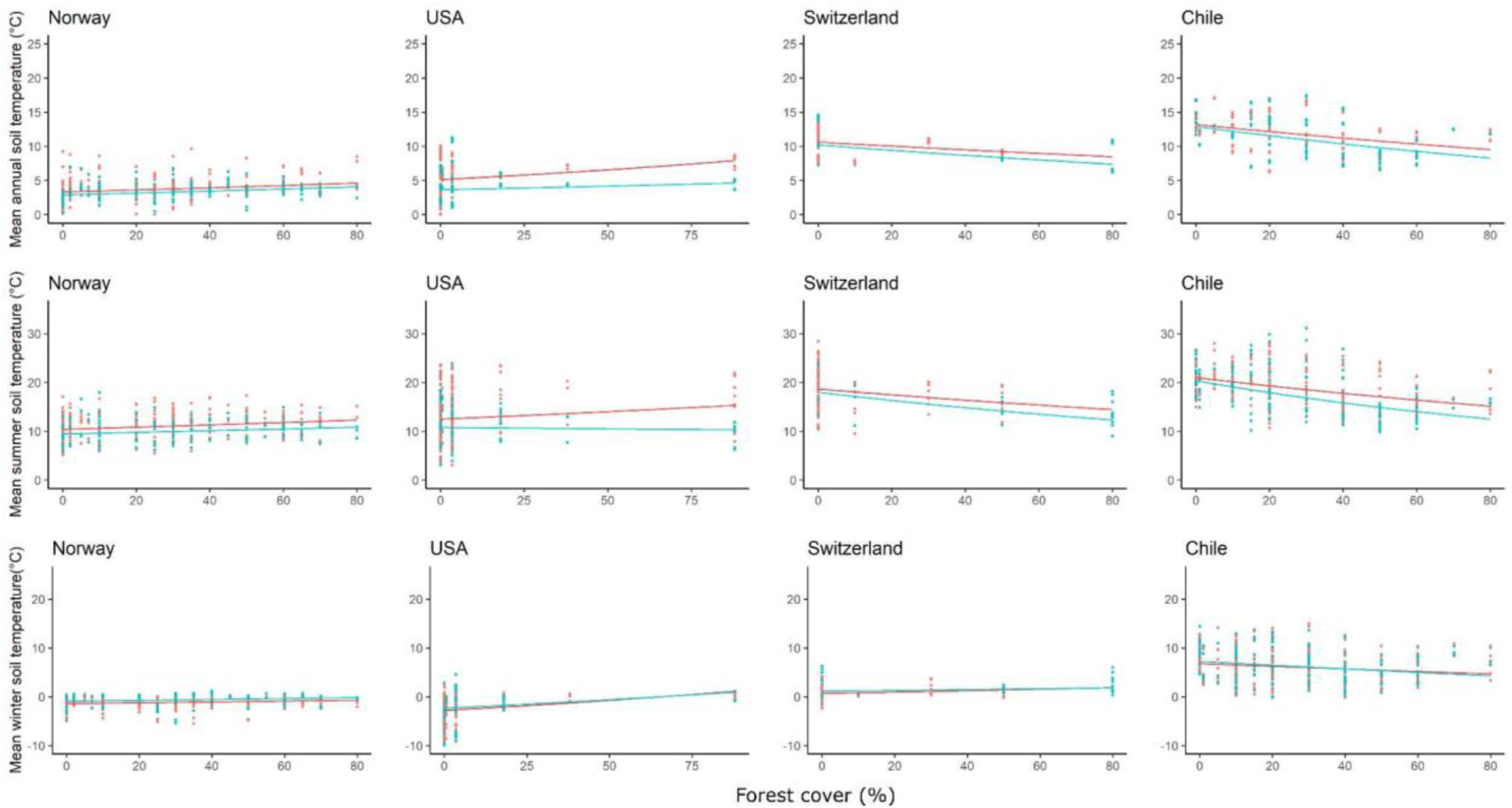
Key soil bioclimatic variables as a function of forest cover (x-axis) and distance to the road (roadside = red, natural vegetation = blue) for four mountain regions. Top row: mean annual soil temperature, middle row: monthly mean soil temperature during summer (July-September on the northern hemisphere, December-February on the southern hemisphere). Bottom row: monthly mean soil temperature during winter (December-February on the northern hemisphere, July-September on the southern hemisphere). Points are raw measurements, fitted curves are from a partial pooling procedure on linear mixed models using the logarithm of the dependent variable (see Table 1). Regions ordered from lowest to highest mean annual temperature in their lowest forest cover plot.

**Figure A2:**
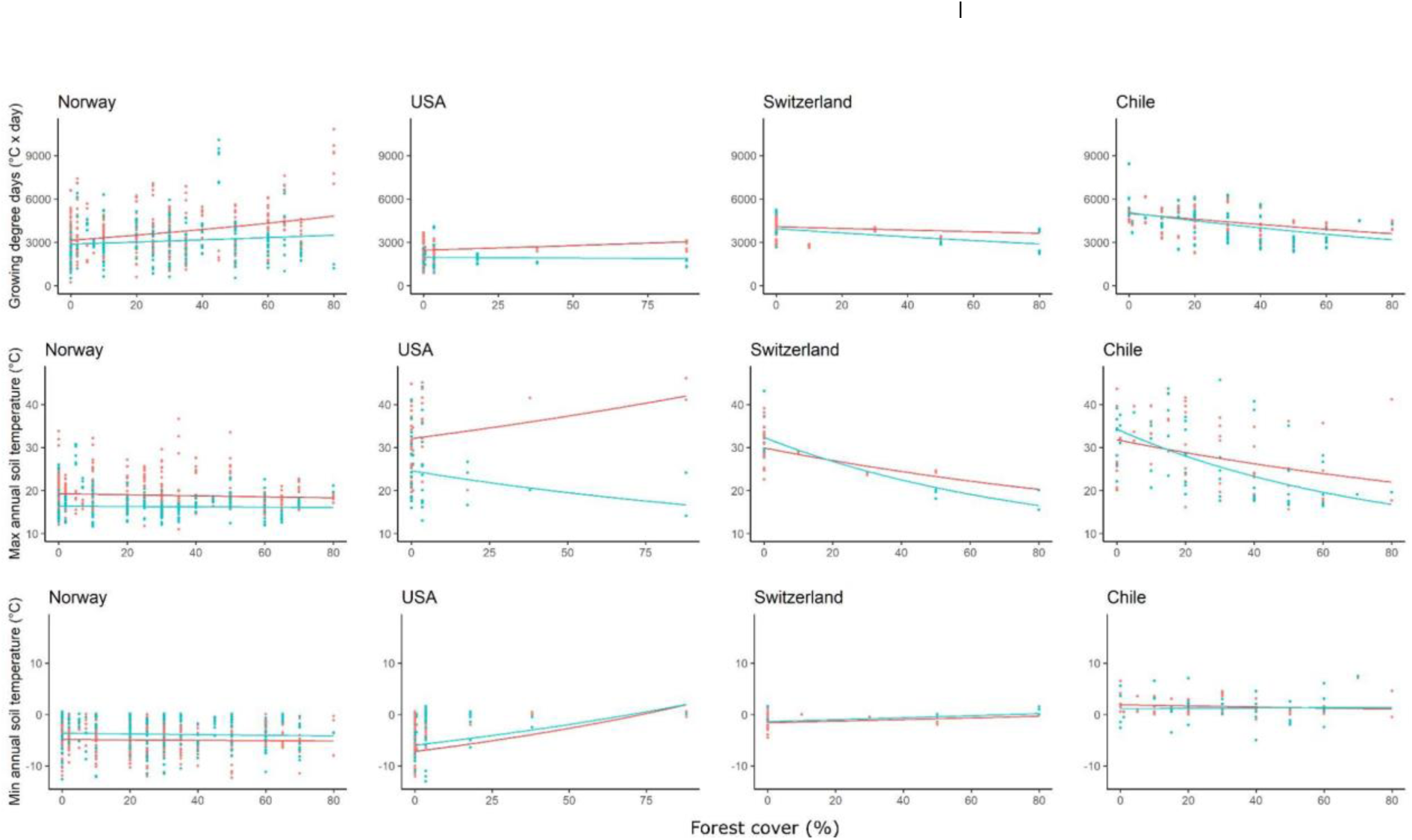
Key soil bioclimatic variables as a function of forest cover (x-axis) and distance to the road (roadside = red, natural vegetation = blue) for four mountain regions. Top row: Growing Degree Days (GDDs), middle row: maximum annual soil temperature. Bottom row: minimum annual soil temperature. Points are raw measurements, fitted curves are from a partial pooling exercise on linear mixed models using the logarithm of the dependent variable (see Table A2). Regions are ordered from lowest to highest mean annual temperature in their lowest forest cover plot.

**Figure A3:**
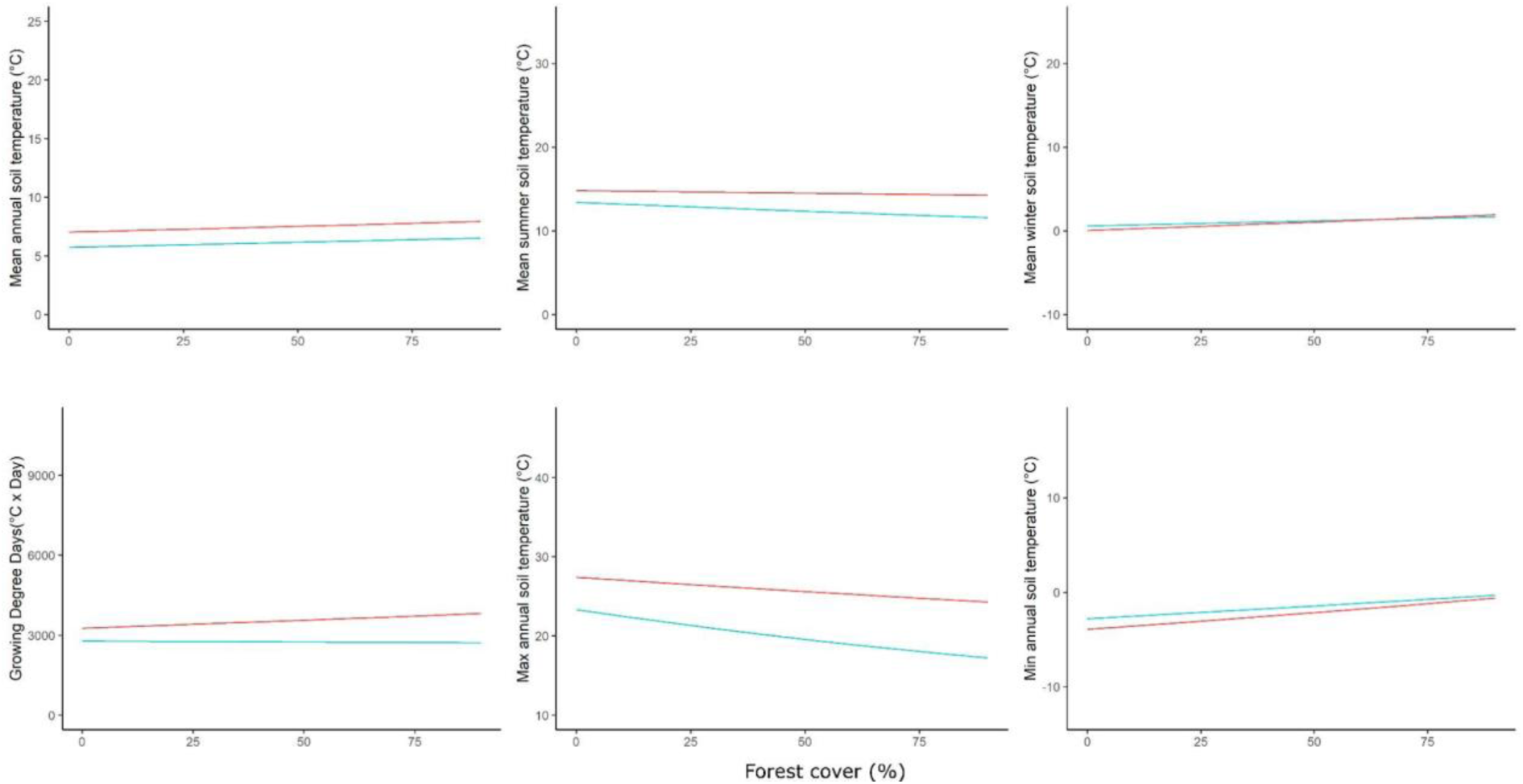
Bioclimatic variables as a function of forest cover (x-axis) and distance to the road (roadside = red, natural vegetation = blue) across all four mountain regions. Top row: mean annual soil temperature, monthly mean soil temperature during summer (July-September on the northern hemisphere, December-February on the southern hemisphere), monthly mean soil temperature during winter (December-February on the northern hemisphere, July-September on the southern hemisphere). Bottom row: Growing Degree Days (GDDs), annual maximum soil temperature and annual minimum soil temperature. Points are raw measurements, fitted curves are drawn using the coefficients of the fixed effects of linear mixed models using the logarithm of the dependent variable.

**Table A1:**
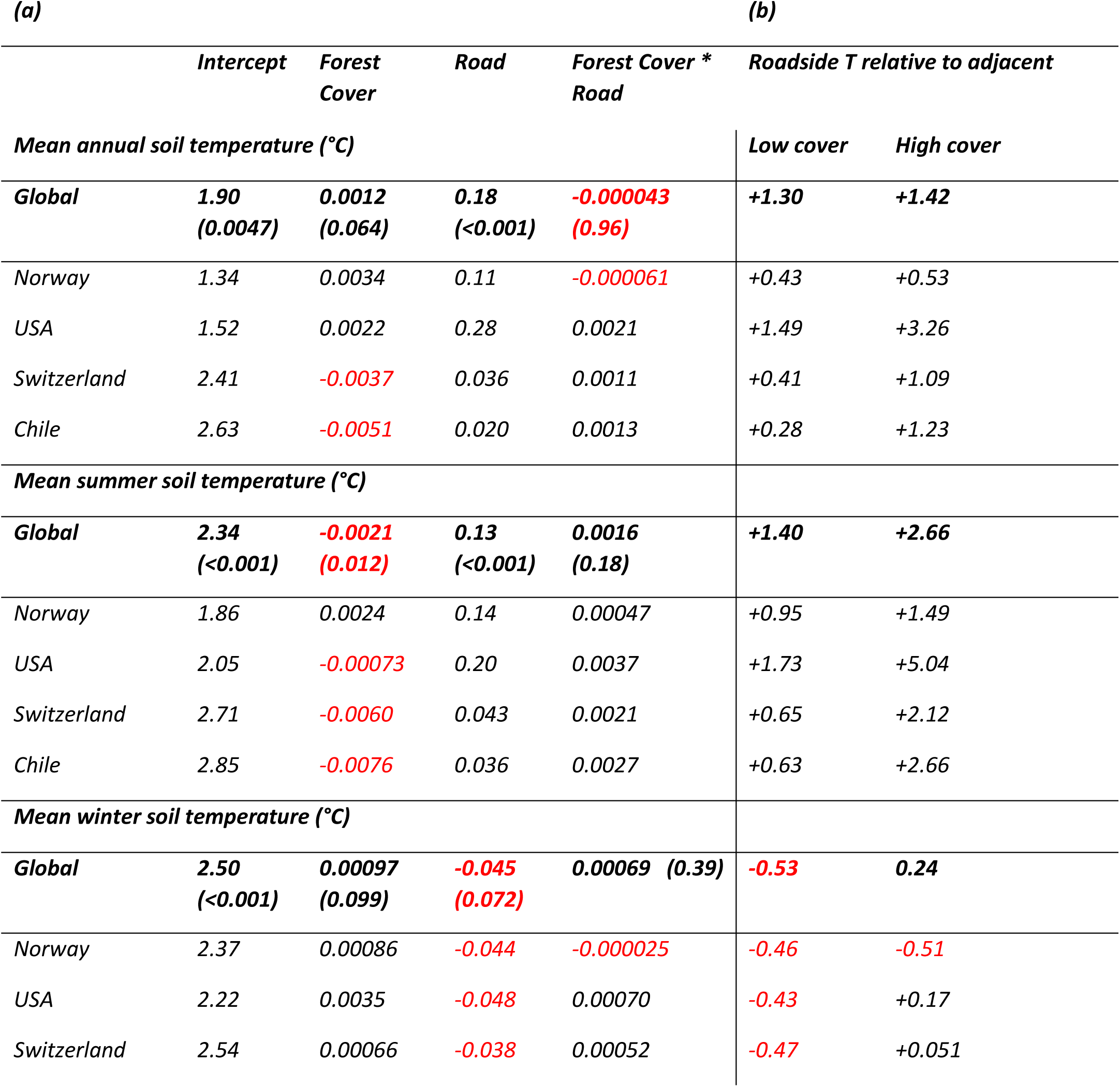
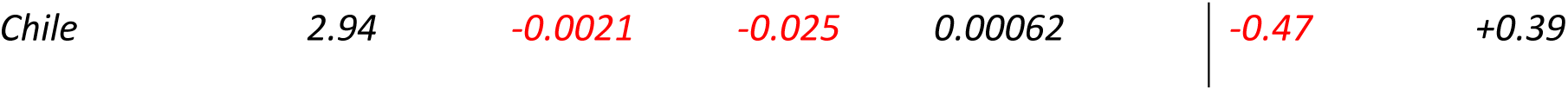
(a) Global coefficients (including p-values) and region-specific coefficients for key soil bioclimatic variables as a function of forest cover and distance to the road for four mountain regions, as in Fig. A1. Top row: mean annual soil temperature, middle row: monthly mean soil temperature during summer (July-September on the northern hemisphere, December-February on the southern hemisphere). Bottom row: monthly mean soil temperature during winter (December-February on the northern hemisphere, July-September on the southern hemisphere). Coefficients are derived from a partial pooling procedure on linear mixed models using the logarithm of the dependent variable. Positive coefficients in black, negative coefficients in red. (b) Modeled temperature differences between the roadside and the adjacent at the lowest and highest coverage percentage of each region’s forest cover gradient. Negative differences (roadsides cooler than adjacent) in red. Regions are ordered from lowest to highest mean annual temperature in their lowest forest cover plot.

**Table A2:**
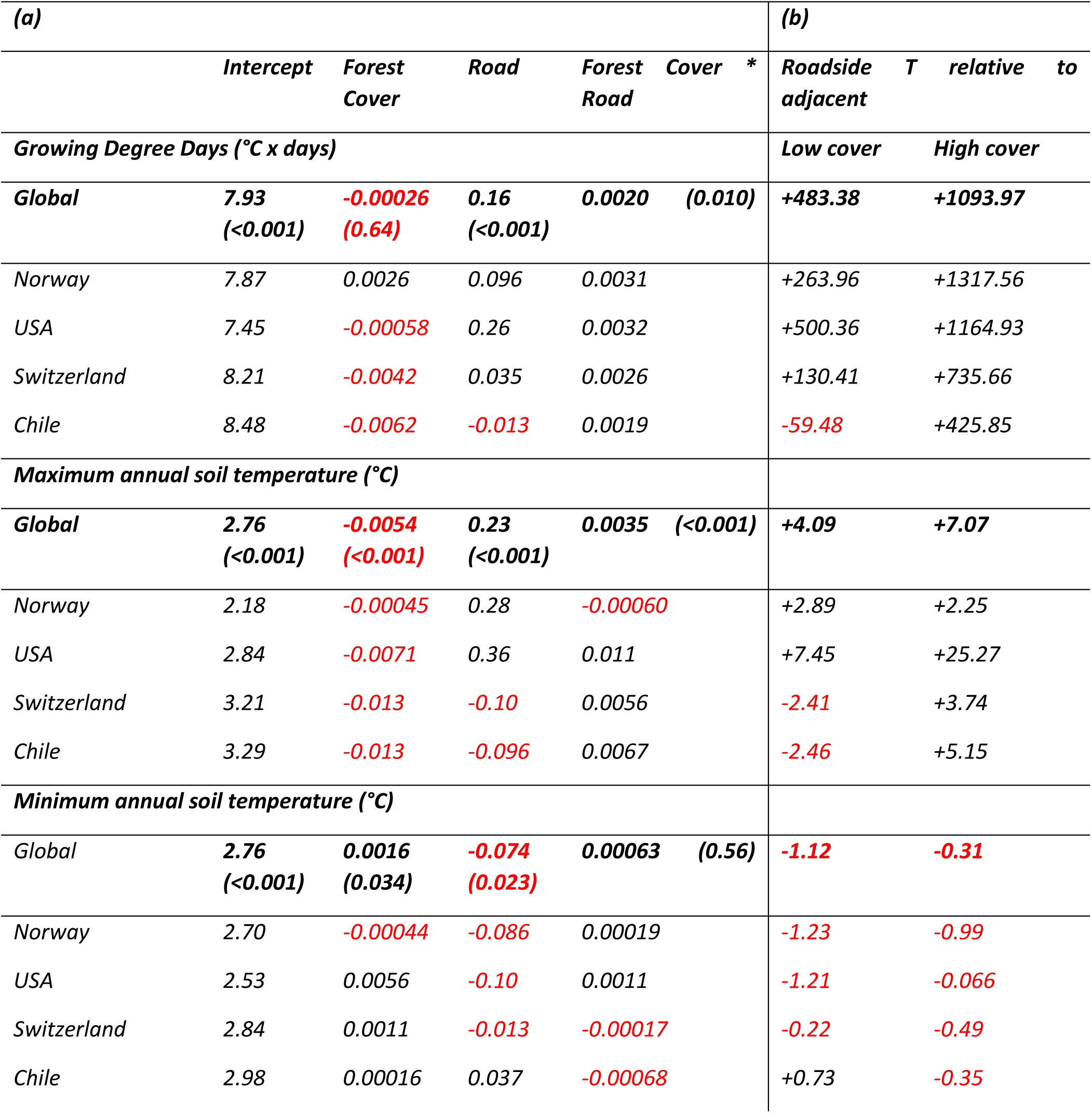
(a) Global coefficients (including p-values) and region-specific coefficients for key soil bioclimatic variables as a function of forest cover (%) and distance to the road (road versus adjacent) for four mountain regions, as in Fig. 2. Top row: Growing Degree Days, middle row: annual maximum soil temperature (95% highest). Bottom row: annual minimum soil temperature (5% lowest). Coefficients are derived from a partial pooling exercise on linear mixed models using the logarithm of the dependent variable. Positive coefficients in black, negative coefficients in red. (b) Modeled temperature differences between the roadside and the adjacent at the lowest and highest coverage percentage of each region’s forest cover gradient. Negative differences (roadsides cooler than adjacent) in red. Regions are ordered from lowest to highest mean annual temperature in their lowest forest cover plot.

